# Plant phenology influences rhizosphere microbial community and is accelerated by serpentine microorganisms in *Plantago erecta*

**DOI:** 10.1101/2021.03.29.437500

**Authors:** Alexandria N. Igwe, Bibi Quasem, Naomi Liu, Rachel L. Vannette

## Abstract

Serpentine soils are drought-prone and rich in heavy metals, and plants growing on serpentine soils host distinct microbial communities that may affect plant survival and phenotype. However, whether the rhizosphere communities of plants from different soil chemistries are initially distinct or diverge over time may help us understand drivers of microbial community structure and function in stressful soils. Here, we test the hypothesis that rhizosphere microbial communities will converge over time (plant development), independent of soil chemistry and microbial source. We grew *Plantago erecta* in serpentine or nonserpentine soil, with serpentine or nonserpentine microbes and tracked plant growth and root phenotypes. We used 16S rRNA barcoding to compare bacterial species composition at seedling, vegetative, early-, and late-flowering phases. Plant phenotype and rhizosphere bacterial communities were mainly structured by soil type, with minor contributions by plant development, microbe source and their interactions. Serpentine microorganisms promoted early flowering in plants on non-serpentine soils. Despite strong effects of soil chemistry, the convergence in bacterial community composition across development demonstrates the importance of the plant-microbe interactions in shaping microbial assembly processes across soil types.

## INTRODUCTION

Plant-microbe associations occur on a small scale, but can impact global patterns, including plant and microbial biodiversity (Cui and He 2009; Ravichandran and Thangavelu 2017; Kandlikar et al. 2019). Plants associate with distinct microbial communities that can benefit plants by enhancing nutrient acquisition (Emami et al. 2018; Fei et al. 2020) and protection against pathogens (De Curtis et al. 2010; Akhtar et al. 2011). These associations generate plant-soil feedbacks which can influence plant community structure (Van Der Heijden et al. 2006). Plant-microbe associations have also been explored for their ability to impact the phenotypes of agricultural plants (Gouda et al. 2018). As a result, microbial amendments are being developed for their ability to influence plant yield (Murgese et al. 2020) and stress tolerance (Orlandini et al. 2014; Kwak et al. 2018). However, the extent to which soil community members establish in the rhizosphere, and when during a plant’s development, remain poorly understood and may affect the efficacy of microbial amendments.

Soil chemistry and plant species both influence the composition of rhizosphere microbial communities (Haichar et al. 2008; Berg and Smalla 2009). Plant development, or phenology, has been shown to correlate with distinct microbial associations. For example, seedlings of *Arabidopsis thaliana* showed distinct microbial communities from later-phase plants (Chaparro et al. 2014). In *Oryza sativa*, rhizosphere microbial communities are dynamic during vegetative growth and can represent particular life phases (Edwards et al. 2018). One mechanism for plant effects on rhizosphere communities is through the rhizodeposition of root exudates, which can change over time and correlates with distinct rhizosphere microbial communities observed at each phase of plant development (Chaparro et al. 2013; Zhalnina et al. 2018). However, whether the same plant species assemble microbial communities similarly in distinct soil backgrounds remains unexplored.

Serpentine soils are characterized by low water-holding-capacity, elevated concentrations of heavy metals, low concentrations of essential plant nutrients, and high Mg to Ca ratios (O’Dell et al. 2006). These characteristics are partially responsible for the low plant productivity and endemism observed on serpentine soils (Anacker 2014). While most plants cannot grow on serpentine soils and other plants can only grow on serpentine soil, serpentine-indifferent plants are able to thrive on serpentine soils and compete on non-serpentine soils (Safford et al. 2005). Serpentine-indifferent plants, with their ability to grow both on and off serpentine soils, are an excellent tool with which to study how soil chemistry influences microbial composition and phenology (Igwe and Vannette 2019). In addition, by utilizing soil treatments with non-adapted microorganisms we can understand how phenology is influenced by different microbial communities.

Abiotic and biotic factors including soil chemistry and soil moisture have also been shown to influence plant phenology; for example, plants growing on serpentine and drought-prone soils have generally been shown to flower sooner than those growing on non-serpentine and non-droughted soils (Sherrard and Maherali 2006; Wright et al. 2006; Rossington et al. 2018; Sakaguchi et al. 2019), but this is not always the case (Schneider 2017). Therefore, an additional goal of our experiments was to investigate the abiotic vs biotic control on phenology, especially as it relates to flowering time.

In this study, we aimed to answer the following broad ecological question: Do plant rhizosphere microbial community grown in disparate soil chemistries converge or diverge over time? More specifically, we test the hypothesis is that soil chemistry influences how microbial communities change over plant development. Further, we predict that rhizosphere microbial communities associated with serpentine-indifferent plants growing on serpentine and non-serpentine soils will become more similar as the plant develops. We also hypothesize that serpentine components introduced to nonserpentine soils, including serpentine microbes, nickel or simulated drought, will change microbial communities and plant characteristics to be similar to those of live serpentine soils. For example, if soil chemistry is the major driver of flowering time, then we can expect that treatments with the same soil origin, regardless of microbial community, will not significantly differ in phenology. If the microbial community influences plant phenology to a greater extent than soil chemistry, we can expect to see significant differences in plant development in soil treatments with non-adapted microorganisms relative to soil treatments with adapted microbes. It is important to understand the relative influence of these factors on plant phenology because the reproductive success of an individual plant and the plant community structure is directly related to plant phenology (Fenner 1998; Rodríguez-Pérez and Traveset 2016; Hidalgo-Triana and Pérez-Latorre 2018).

## METHODS

### Study system and soil collection

Soils were collected from McLaughlin Natural Reserve in June 2018 from three serpentine and three nonserpentine sites (Sites 1, 2, and 3 from Igwe and Vannette 2019). McLaughlin Natural Reserve is characterized by a Mediterranean climate with hot and dry summers from April to October. Gallon-sized plastic bags of soil were collected every 5-meters across a 20-meter transect at each site at an average depth of 10 cm. These soils were placed on ice in the field and then in Sterilite plastic containers at 4°C until the start of the experiment in July 2018. We used *Plantago erecta* (Serpentine Affinity Mean = 1.0), which is common in serpentine and non-serpentine sites locally (Safford et al. 2005). Seeds used in the experiment were purchased from S&S Seeds in 2016 (Carpinteria, CA) after field-collected seeds germinated poorly.

### Growth chamber experiment

We conducted an experiments aimed to examine how background soil chemistry and soil microbial community jointly influence plant growth and microbial community assembly in the rhizosphere. All soils included an autoclaved soil background (serpentine or nonserpentine), to which live (unautoclaved) soils from either serpentine or nonserpentine soils were added at approximately 16% (v/w) to create the microbial amendments (Farrer and Suding 2016; Calderón et al. 2017). Soils were autoclaved at 120°C at 15 psi for two 30-min periods with 24-hours between sterilizations (Ishaq et al. 2017). Using this method, four factorial treatments were created: autoclaved serpentine soil with serpentine microbes (S+Sm), autoclaved nonserpentine soil with nonserpentine microbes (NS+NSm), autoclaved serpentine soil with nonserpentine microbes (S+NSm), autoclaved nonserpentine soil with serpentine microbes (NS+Sm). In addition, to explore which dimensions of serpentine soils shape plant-microbe-soil interactions (Wright et al. 2006), we amended some NS+Sm treatments with either nickel (final concentration of 25 ppm; NS+Sm+Ni), or grew plants in conditions simulating drought stress (NS+Sm+Drought).

*Plantago erecta* seeds were added to D16 Deeppots (volume: 16 in^3^ and 262 mL; Stuewe and Sons., Inc, Tangent, OR) containing ∼100 g of soil in one of six soil chemistries above. Plants were grown in a growth chamber under 12:12 light/dark regime at 20°C at the UC Davis Environmental Horticulture Greenhouse Complex and were grown to senescence, with 15 replicates per treatment and 3 non-planted controls per treatment. The simulated drought soil treatment was watered until the soil was saturated once a week while all other soil treatments were watered daily with DI water. Leaf number and plant height were recorded weekly until senescence. Plants in each treatment were harvested at seedling, vegetative, early flowering, and late flowering phases and a random subset (N=6) were used for rhizosphere soil collection, microbial DNA extraction, and 16S rRNA sequencing. Plants were classified as ‘seedlings’ upon emergence from the soil. When true leaves were present, plants were classified as ‘vegetative’. ‘Early flowering’ was characterized by shoot development and the presence of an undeveloped terminal protuberance. Once the plant began to bloom, the plant was characterized as ‘late flowering’. Once the plant became brittle to the touch, they were classified as ‘senescing’. For two treatments (NS+Sm+Ni) and (NS+Sm+Drought), only seedling and late flowering phases were harvested due to limitation in growth chamber space.

### Rhizosphere soil collection

For each harvested plant, roots were shaken to remove loosely adhering soil. An ethanol-sterilized razor was used to separate the stem from the roots. Above-ground plants were dried at 80°C for 48 hours and then weighed.

Roots were separated from rhizosphere soils. Briefly, roots were sonicated in 0.9% NaCl/0.01% Tween 80 (v/v) solution for 180 seconds to remove the tightly adhering soil particles (Barillot et al. 2013). Centrifuge tubes containing NaCl/Tween (without roots) were then centrifuged for 20 minutes at 4°C at 3234 x g. The pellet was frozen at -20°C until DNA extraction using a Zymo fecal/soil DNA extraction kit according to enclosed directions (Zymo Research, Irvine, CA). After sonication, root samples were stored in 50% ethanol solution until root image analysis. DNA extraction was confirmed using a Nanodrop 1000 spectrophotometer (ThermoScientific, Waltham, MA, USA) then samples were submitted to the Centre for Comparative Genomics and Evolutionary Bioinformatics Integrated Microbiome Resource at Dalhousie University for PCR and sequencing. The V6-V8 subregion of the 16S SSU rRNA was amplified using B969F (ACGCGHNRAACCTTACC) and BA1406R (ACGGGCRGTGWGTRCAA) primers (Comeau et al. 2011). DNA was amplified using Phusion High-Fidelity DNA polymerase (NEB) and MiSeq (300+300 bp PE) for final amplicon lengths that were 508 bp. Raw sequences are archived at www.ncbi.nlm.nih.gov/sra/PRJNA623253.

### Root imaging

To analyze root length, volume, surface area and diameter, samples were scanned using WinRHIZO optical scanner and software (Regent Instruments Inc., Canada). Each root sample was imaged individually by laying them flat onto a tray containing 50% ethanol to cover the entire root. Tangled roots were carefully separated with forceps and any roots broken off were also imaged. Before root parameters were measured, any residual soil particles and foreign root fragments scanned by the imaging software were eliminated from the selected root image. To do this, the entire root was selected by drawing a box around the imaged root. Next, foreign particles were excluded from analysis by selecting *Regions - Exclusion Regions* and drawing a box around each individual target region. Once this step was completed, root parameters including root length, root diameter, root surface area, and root volume were measured by selecting *Image - Image with Analysis*.

### Bioinformatics

Amplicon sequence variants (ASVs) from 16S rRNA amplicons were identified using DADA2 (v1.7.2) (Callahan et al. 2016a). Briefly, paired-end fastq files were processed by filtering and truncating forward reads at position 250 and reverse reads at position 200. Sequences were dereplicated, merged and error-corrected according to code archived on Dryad. Chimeras were removed, and the taxonomy assigned using the SILVA database (v128) (Quast et al. 2012; Yilmaz et al. 2014; Glöckner et al. 2017). A phylogenetic tree based on 16S sequences was created using the DECIPHER package (v2.8.1) in R to perform multi-step alignment and phangorn (v2.4.0) to construct the tree using neighbor-joining (Wright et al. 2006; Schliep 2011). The sequence table, taxonomy, and metadata were combined into a phyloseq object and used for further analysis (phyloseq v1.30.0) (McMurdie and Holmes 2013; Callahan et al. 2016b). Mitochondrial and chloroplast sequences as well as any sequences that were not assigned to bacteria were removed from the ASV table.

### Statistical analysis

To visualize the relative abundance of each phylum, ASVs were aggregated to the phylum level and taxa representing less than 2% of relative abundance were filtered out. To determine effect of soil treatment and plant developmental phase on alpha diversity of rhizosphere communities, Shannon diversity was calculated on the full dataset using the estimate_richness function in the phyloseq package (1.30.0) and used as a response variable in ANOVA with plant developmental phase, soil chemistry (S, NS, NS+Sm+Ni, and NS+Sm+D), and microbe source (S or NS) as predictors. Shannon index was used because it accounts for both abundance and evenness in samples (Kaisermann et al. 2017).

To examine differences in rhizosphere bacterial species composition due to soil chemistry, microbe source, and plant developmental phases, Bray-Curtis dissimilarities were calculated and visualized using non-metric multidimensional analysis (NMDS). To determine which predictors were associated with variation in rhizosphere bacterial composition, we used the ‘adonis’ function from the vegan package with Bray-Curtis dissimilarities as the response variable and plant developmental phase, soil chemistry, and microbe source as predictors. To test for differences in multivariate dispersion among rhizosphere communities, the ‘betadisper’ function from the vegan (v2.5.3) package was used (Oksanen et al. 2019) with soil chemistry, plant developmental phase, and microbe source as predictors.

To determine the effects of soil treatments on plant growth, each plant trait (leaf number, plant height, root length, root diameter, root surface area, and root volume), was analyzed using a general linearized model with soil chemistry, microbe source, and plant developmental phase as the predictor and differences between group means were identified using likelihood ratio tests. Tukey HSD was used as a post-hoc test to identify differences among groups.

To determine the effects of soil treatments on time to flowering, survival analysis was conducted on binary flowering data (yes/no) using Kaplan-Meier curves and a Cox proportional hazards regression model (‘coxph’) on a survival object (‘Surv’) in the survival package (v2.42.6) to describe the soil treatment impacts the probability of flowering over time (Therneau 2015). The time to event (late flowering), was measured in days from the onset of seedling phase. The model provides a hazard ratio (HR) where a HR > 1 indicates an increased likelihood of development, while an HR < 1 indicates a decreased likelihood of development. Differences in the hazard ratio were visualized using ‘ggforest’ in the survminer package (v0.4.3) (Kassambara and Kosinski 2018).

To determine if the relative abundance of bacteria ASVs differed among plant developmental phases or soil chemistries, differential abundance analysis using DESeq2 (1.26.0) was used with soil chemistry as the predictor. DESeq2 analysis was conducted with all soil treatments for the seedling and late flowering plant developmental phase and another analysis was conducted with S+Sm, NS+NSm, S+NSm, and NS+Sm for all plant developmental phases to represent the experimental design.

## RESULTS

After quality filtering and removal of non-target sequences, we recovered 1,608,688 reads (average 9,353 reads per sample) that were grouped into 23,894 amplicon sequence variants. Sampling curves within most samples were saturating (NS+NSm, NS+Sm+Ni, NS+Sm+D) indicating a robust sampling of the microbial diversity associated with individual plants while serpentine soils did not fully saturate (S+Sm, S+NSm) (Supplemental Figures 1-6). To compare among samples, count data were normalized by relative abundance in a sample.

### Soil treatment and plant developmental phase influence species richness and community similarity

Bacteria from the phyla Proteobacteria or Acidobacteria comprised nearly 90% of reads from most samples depending on soil chemistry (Figure 1a). Alpha diversity differed among soil chemistries (Figure 1b; F_3,100_ = 124.05, *P* <0.001) and plant developmental phase (F_4,100_ = 5.74, *P* = <0.001), but not microbe source (F_1,100_ = 2.64, *P* = 0.107). There was a significant interaction between soil chemistry and microbe source (F_1,100_ = 6.80, *P* = 0.011) as well as soil chemistry, microbe source, and plant developmental phase (F_5,100_ = 4.08, *P* = 0.004) on bacterial composition. The serpentine soil treatments (S+Sm and S+NSm) had lower alpha diversity at all time points compared to all treatments with nonserpentine soils (NS+Sm, NS+Sm+Ni, NS+Sm+D). The treatment with simulated drought (NS+Sm+D) treatment had higher species richness than either the live nonserpentine treatment (NS+NSm) or the treatment with nickel added (NS+Sm+Ni).

**Figure 1.**
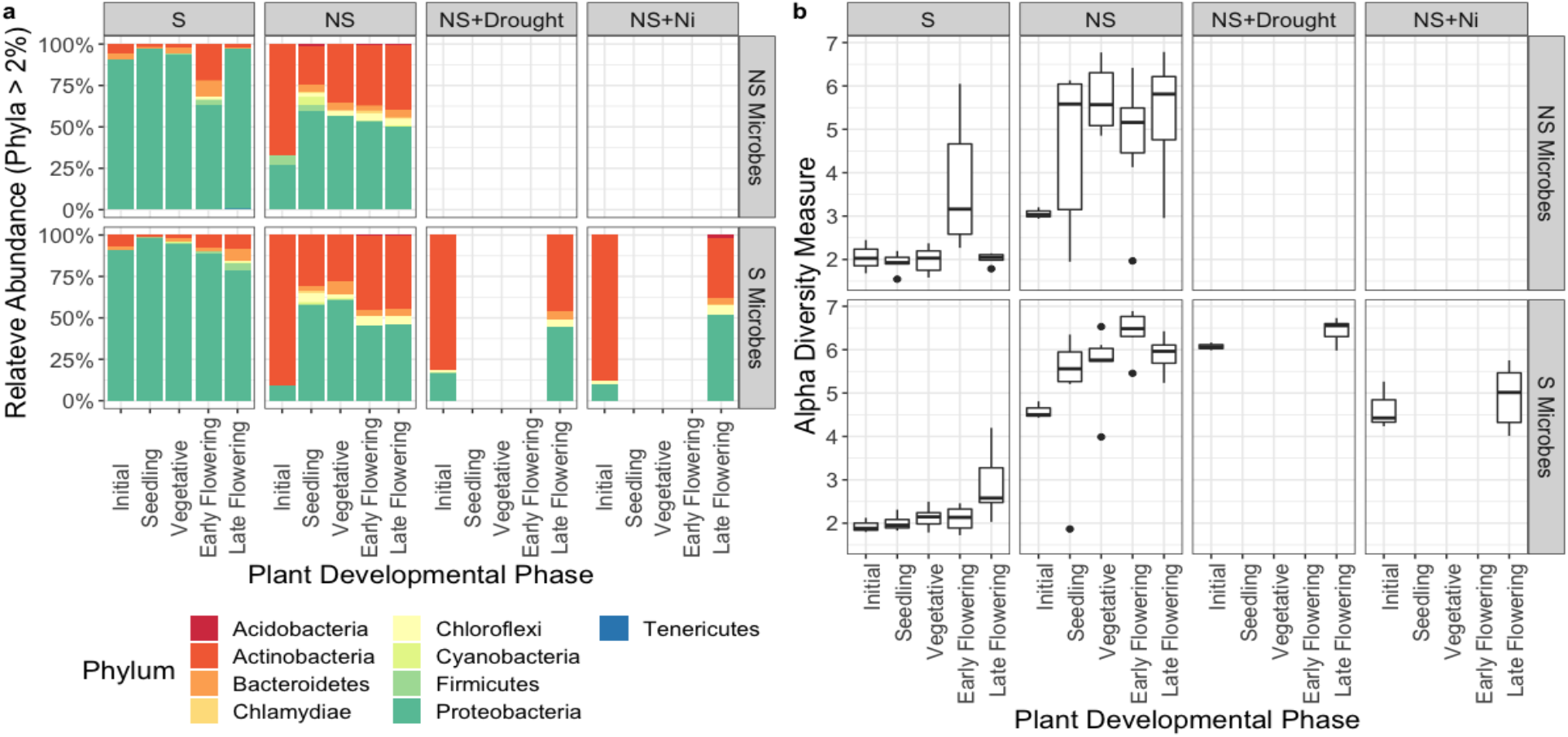
Relative abundance and alpha diversity of bacterial community composition on roots of *Plantago erecta* across soil treatments and plant phenology. (a) Bacterial phyla with a relative abundance of at least 2% are visualized in a bar graph facetted by plant developmental phase and soil treatment (S = Serpentine and NS = Nonserpentine). (b) Shannon diversity is significantly different between soil chemistries (F_3,100_ = 124.05, P <0.001) and plant developmental phase (F_4,100_ = 5.74, P <0.001), but not microbe source (F_1,100_ = 2.64, P = 0.11). There was a significant interaction between soil chemistry and microbe source (F_1,100_ = 6.80, P = 0.011) as well as soil chemistry, microbe source, and plant developmental phase (F_5,100_ = 4.08, P = 0.004).

Bacterial community composition in the rhizosphere varied with soil chemistry and plant developmental phase, with surprisingly minimal contribution from the microbial source (Figure 2; Table 1). Variability in bacterial communities among plants (beta diversity) was associated with soil chemistry (Betadisper: F_3,128_ = 93.67, *P* = 0.001) and plant developmental phases (Betadisper: F_4,127_ = 5.79, *P* = 0.004), but only weakly with microbe source (Betadisper: F_1,130_ = 3.42, *P* = 0.068). Microbial communities from the seedling phase were less variable than all other plant developmental phases, but there were no significant differences between the variability of other phases. Both serpentine soil treatments were less variable than that of either nonserpentine soil treatments.

**Table 1.** Statistical analysis of microbial community dissimilarity using ANOVA. The predictors, degrees of freedom (df), number of samples (N), F-Value (F), Variation, and P-Value are listed

| Predictor | df | N | F | R<sup>2</sup> | P |
| --- | --- | --- | --- | --- | --- |
| Soil type | 3 | 100 | 22.23 | 0.304 | 0.001 |
| Developmental phase | 4 | 100 | 3.21 | 0.059 | 0.001 |
| Microbe source | 1 | 100 | 3.24 | 0.015 | 0.008 |
| Soil type*Developmental phase | 6 | 100 | 2.21 | 0.06 | 0.001 |
| Microbe source*Developmental phase | 4 | 100 | 1.36 | 0.025 | 0.089 |
| Soil type*Microbe source | 1 | 100 | 3.04 | 0.014 | 0.011 |
| Soil type*Microbe source*Developmental phase | 4 | 100 | 1.58 | 0.029 | 0.031 |

**Figure 2.**
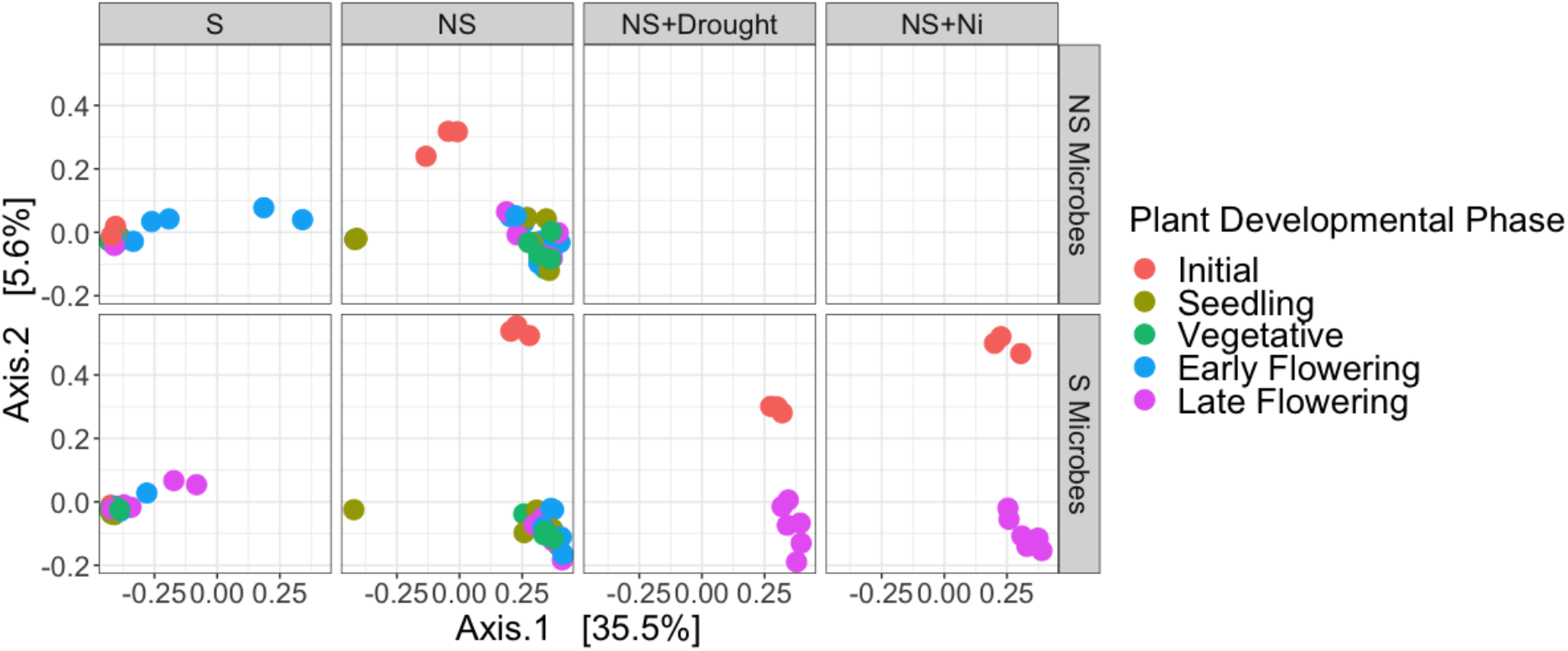
PCoA of *Plantago erecta* rhizosphere bacterial communities across soil treatments and plant phenology using Bray-Curtis dissimilarity. Point color indicates plant developmental phase and panels indicate distinct soil treatments.

### Plant growth responses to soil chemistry and microbial source

In general, the height, leaf number, aboveground dried biomass, root length, root diameter, root surface area, and root volume were all impacted by soil chemistries (Figures 3 and 4; Table 2 and 3). The interaction between plant developmental phase and microbe source as well as soil chemistry and microbe source also influenced these plant traits. Microbe source, alone, only significantly influenced root diameter.

**Table 2.** Statistical results for aboveground plant metrics. Linear mixed-effects model was used to determine the impact of various predictors on plant height, leaf number, and dry biomass. The tray where plants were grown was used as a random variable.

| Predictor | df | N | Plant Height |  | Leaf Number |  | Dry Biomass |  |
| --- | --- | --- | --- | --- | --- | --- | --- | --- |
|  |  |  | X2 | P | X2 | P | X2 | P |
| Soil chemistry | 3 | 3372 | 4890.83 | <0.0001 | 78.44 | <0.0001 | 60.40 | <0.0001 |
| Developmental phase | 3 | 3372 | 7012.10 | <0.0001 | 3064.16 | <0.0001 | 79.84 | <0.0001 |
| Microbe source | 1 | 3372 | 2.58 | 0.108 | 137.12 | <0.0001 | 2.45 | 0.11784 |
| Soil chemistry*Developmental phase | 9 | 3372 | 1294.27 | <0.0001 | 57.13 | <0.0001 | 35.69 | <0.0001 |
| Microbe source*Developmental phase | 3 | 3372 | 40.84 | <0.0001 | 20.45 | 0.0001 | 8.81 | 0.03193 |
| Soil chemistry*Microbe source | 1 | 3372 | 296.64 | <0.0001 | 142.12 | <0.0001 | 2.34 | 0.12583 |
| Soil chemistry*Microbe source*Developmental phase | 3 | 3372 | 11.79 | 0.008 | 53.86 | <0.0001 | 6.72 | 0.08134 |

**Table 3.**
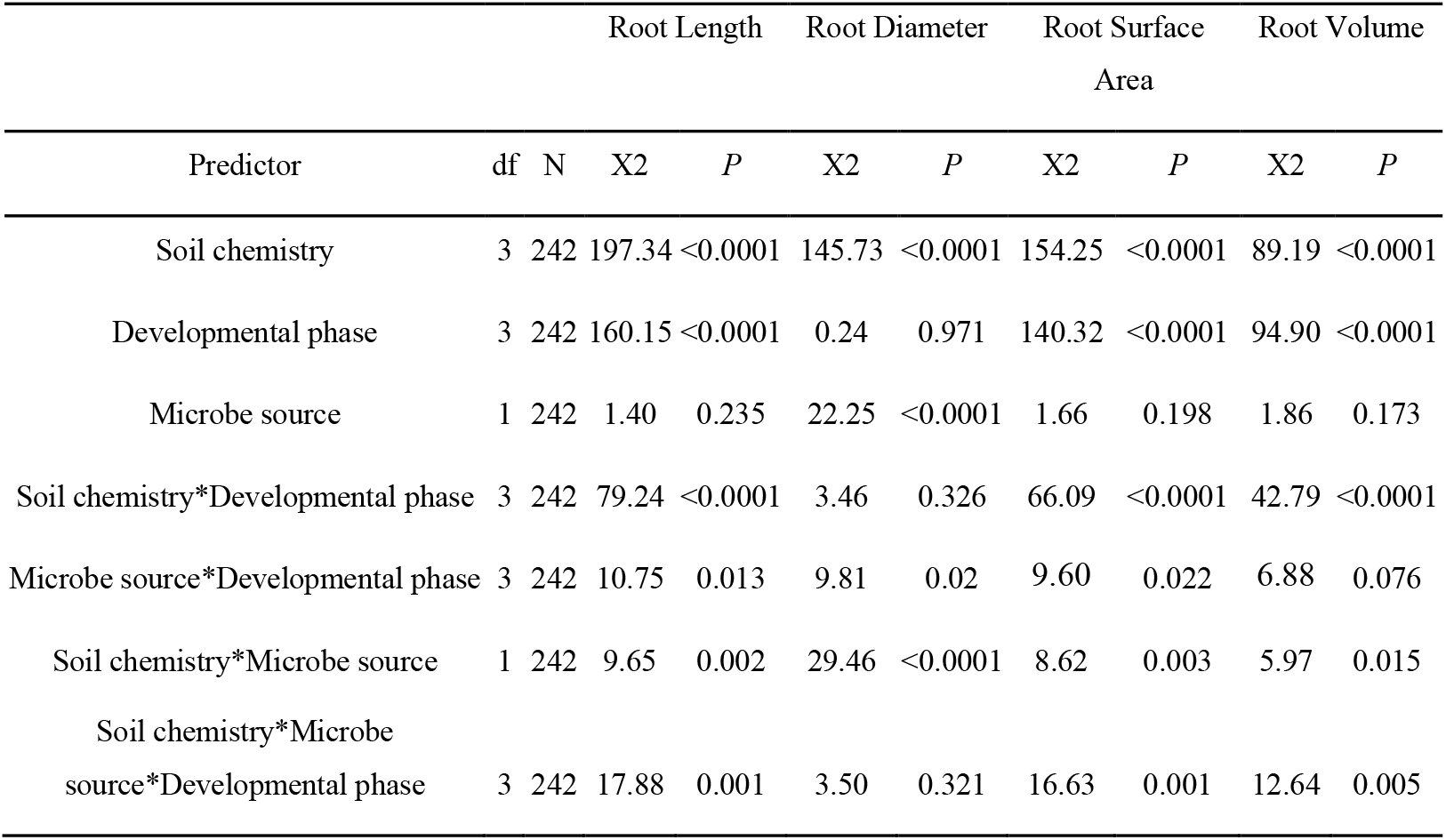
Statistical results for belowground plant metrics. Linear mixed-effects model was used to determine the impact of various predictors on root length, diameter, surface area, and volume. The tray where plants were grown was used as a random variable.

**Figure 3.**
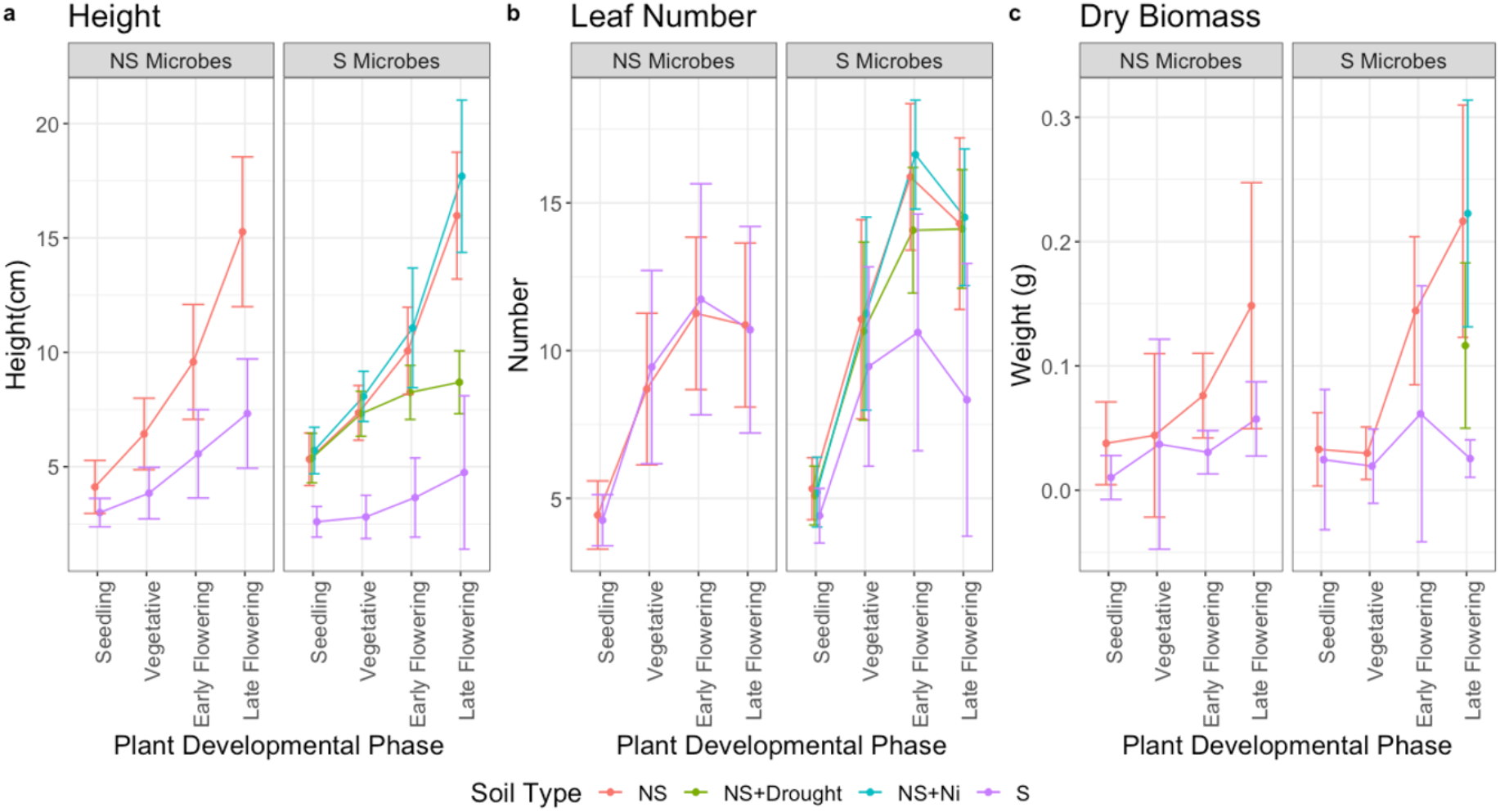
Growth traits of *Plantago erecta* vary among soil treatments and plant developmental phase. Points indicate mean +/- 1SD and points from soil treatments are connected, showing there is a significant difference between (a) height and (b) leaf number between soil treatment and plant developmental phase. Dry biomass (c) was significantly different between plant developmental phase, but not soil treatment.

**Figure 4.**
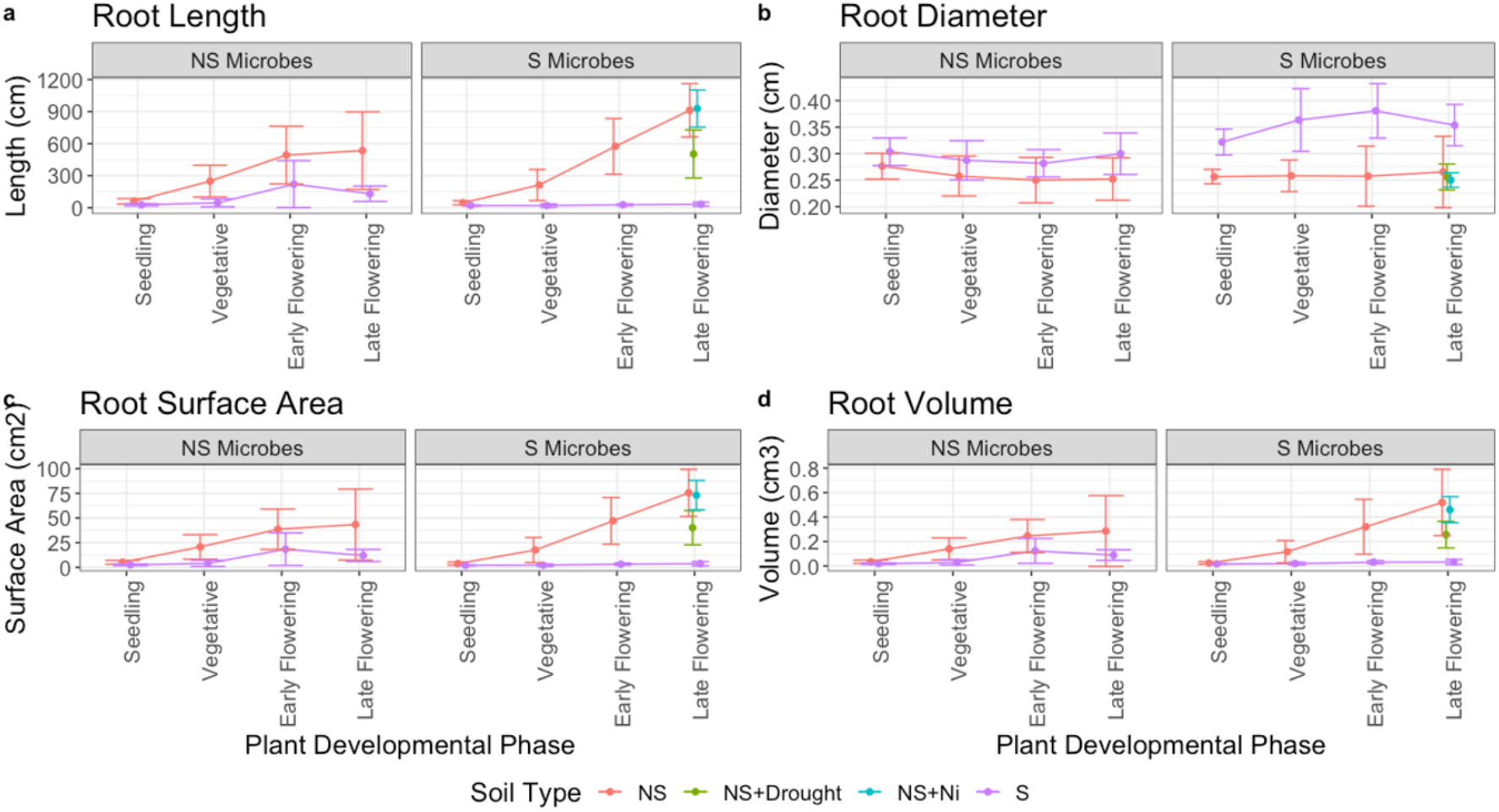
Root growth traits vary among soil treatments and plant developmental phase. Mean value for root traits and standard deviation show that (a) root length was significantly different across soil treatment and plant developmental phase. Root diameter (b) was different between soil treatments, but not plant developmental phase. Root surface area (c) and root volume (d) showed significant differences between plant developmental phases and soil treatments. The interaction between soil treatment and plant developmental phase influenced all root metrics.

### Serpentine microbes alter plant vegetative and flowering phenology

Cox proportional hazards regression models showed differences in plant progression through developmental phases among treatments (Table 4). Plants associated with serpentine microbes reached the post-seedling vegetative phase and flowered earlier than those associated with nonserpentine microorganisms (Table 4).

**Table 4.** Cox proportional hazards model to determine differences in *Plantago erecta* phenology. Cox proportional hazards model was used to determine the likelihood of *P. erecta* reaching a particular plant development phase in distinct soil types. A HR=1 indicates the treatment was used as a reference to which other treatments were compared. A HR > 1 indicates an increased likelihood of development, while an HR < 1 indicates a decreased likelihood of development. For example, *P. erecta* grown in Serp+NSmic is 3.1 times more likely to reach the flowering phase than those grown in live serpentine soil (Serp). Values in parentheses are the confidence intervals for the hazard ratio and * indicates p-value ≤ 0.001.

| Soil Type | Plant Developmental Phase |  |  |
| --- | --- | --- | --- |
|  | Vegetative | Early Flowering | Late Flowering |
| S+Sm | 1 | 1 | 1 |
| NS+NSm | 1.38<br>(0.94-2.0) | 1.7<br>(0.99-3.1) | 1.3<br>(0.68-2.6) |
| S+NSm | 0.84<br>(0.56-1.2) | 1.2<br>(0.65-2.1) | 1<br>(0.52-2.1) |
| NS+Sm | 6.23*<br>(4.03-9.6) | 4.2*<br>(2.36-7.6) | 3.1*<br>(1.56-6.1) |
| NS+Sm+Ni | 7.45*<br>(4.02-13.8) | 6.3*<br>(3.02-16.1) | 2.1<br>(0.99-4.5) |
| NS+Sm+D | 4.74*<br>(2.61-8.6) | 9.9*<br>(4.78-20.3) | 30.9*<br>(11.52-83.0) |

### Bacterial taxa

DESeq2 identified taxa that were differentially abundant according to soil type (Figure 5, Supplemental Figure 7-8). *Solirubrobacter, Lactobacillus*, and *Methylobacterium* were genera that were of particular interest. *Solirubrobacter* were present in all treatments with nonserpentine soils and absent in both treatments with serpentine soils. *Lactobacillus* were present in both serpentine soil treatments. *Methylobacterium* were most abundant in the NS+NSm, NS+Sm, and S+Sm treatments.

**Figure 5.**
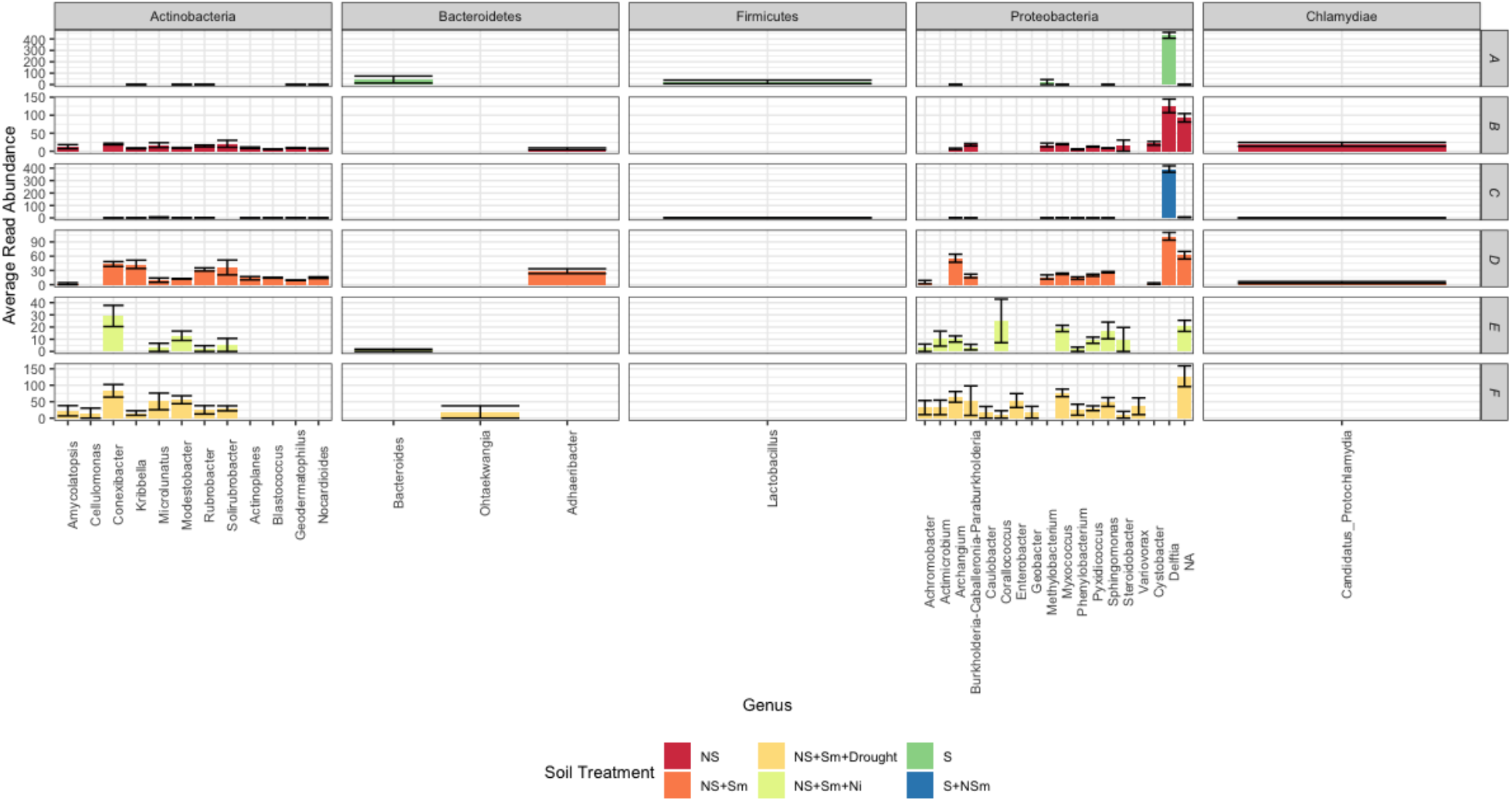
Differentially abundant genera across soil treatments. DESEq2 analysis showing ASVs that were differentially abundant between soil treatments (FDR <0.01). Bacterial genus is on the x-axis and relative average read abundance on the y-axis. Colors represent soil treatments (A=Serpentine soil and serpentine microbes, B=Nonserpentine soil and nonserpentine microbes, C=Serpentine soil and nonserpentine microbes, D=Nonserpentine soil and serpentine microbes, E=Nonserpentine soil and serpentine microbes and nickel stress, F=Nonserpentine soil and serpentine microbes and drought stress). Bars represent means +/- 1SE.

## DISCUSSION

The role of rhizosphere microbes in plant health is increasingly recognized, but efforts to manage or alter rhizosphere composition require understanding the relative importance of soil chemistry, microbial species pools and plant development in assembly processes. In comparing serpentine to nonserpentine soils and microbial sources, we found that soil chemistry exerts that strongest influence on microbial community composition, with more minor changes with plant phenology and with microbial source. Plant phenology was also impacted by soil chemistry and microbe source with plants growing in serpentine soils having a delayed vegetative and flowering phenology. Changes in flowering phenology can have an impact on life-history traits and population dynamics (Dorji et al. 2013; Yang et al. 2020).

### Soil chemistry and plant developmental phase both significantly impact microbial diversity and community composition

Here, rhizosphere alpha diversity generally increased with plant developmental phase and was generally higher when plants were grown in nonserpentine soils, which allowed plants to grow larger. Previous research found no or minimal difference in bacterial alpha diversity between serpentine and nonserpentine soils (Oline 2006; Igwe and Vannette 2019). Consistent with our previous research, plants grown on nonserpentine soils showed increased alpha diversity, suggesting that plant growth rather than the diversity of microbes in the species pool is more important in determining bacterial diversity in the rhizosphere. This may be due to accumulation of microbes simply due to the amount of time the plants spent in soil (Dombrowski et al. 2017) or changes in the amount or type of exudates deposited in the rhizosphere (Chaparro et al. 2013; Zhalnina et al. 2018).

The largest change between microbial composition occurred between the start of the experiment and the seedling phase. After the seedling phase, the microbial community composition stabilized. This occurrence is in line with previous research in rice that showed similar results (Edwards et al. 2018). Some microbial communities in the serpentine soil treatments at the flowering phases shift to look more similar to those associated with nonserpentine *P. erecta*. Therefore, some convergence is occurring; however, in this experiment, soil chemistry contributed more to the observed beta diversity in the microbial community than plant developmental phase, even at later phases.

Various mechanisms could contribute to the observed results. For example, the presence of DNA from dead cells that saturated sequencing efforts relative to new DNA. The soil could also exert a selective pressure that is stronger than that of *P. erecta* rhizosphere. Alternatively, is possible that some serpentine microbes that grow well in nonserpentine soils and vice versa. It has been shown that serpentine and nonserpentine soils can host the same microbes with varying accessory genomes (Porter et al. 2017). Shotgun sequencing or whole genome sequencing could identify if the same microbes with distinct genotypes grew in reciprocal soil chemistries. By exuding carbon compounds, phenolic acids, and amino acids, plants can enhance the growth of specific beneficial or pathogenic members of the soil microbial community, which can enhance plant growth in some cases (Paterson et al. 2007). Root exudates change over the course of plant development where younger plants exude more sugars while older plants exude more complex carbon compounds. Characterizing root exudation of *P. erecta* over plant development and correlating the results with changes in microbial community composition can provide greater insight into the role of dynamic plant exudation on survival in serpentine and nonserpentine soils.

Our study cannot disentangle the possible mechanisms that contribute to the observed results. We sampled plants at distinct development phases irrespective of soil residence time which is an experimental design that considers that plant development phase was shown to influence the rhizosphere microbial community separately from chronological age (Edwards et al. 2018). In addition, our study was performed in a growth chamber, preventing the opportunity for microbial immigration from the soil, which could affect microbial diversity and composition in the field.

### Soil microbial community and soil chemistry influence time to vegetative growth and flowering

Our reciprocal transplants revealed that serpentine microorganisms, when in non-serpentine soils, accelerate vegetative, early flowering, and flowering phenology (Table 4). The importance of the microbial community for time to flowering has been previously demonstrated in *Arabidopsis* (Wagner et al. 2014) and *Ipomea purpurea* (Chaney and Baucom 2020). Drought-adapted microbes accelerated flowering in *Brassica* when compared to non-drought-adapted microbes (Lau and Lennon 2012). A few mechanisms for microbial effects on phenology have been proposed including nutrient availability, production of plant hormones or their precursors, or by exacerbating stress. However, flowering was delayed in the presence of serpentine microbes grown in nonserpentine soils (Table 4), which have been previously demonstrated to be more nutrient-rich than serpentine soils (Brady et al. 2005), suggesting another mechanism may underlie microbial effects in this experiment. It may be that microbes produce plant hormones such as indole acetic acid (IAA) which plays a significant role in flowering time. Nitrogen can be converted to tryptophan and then to IAA and increases time to flowering (Lu et al. 2018). Alternatively, flowering time has been shown to be impacted by biotic stress (Kazan and Lyons 2016). It is possible that introducing non-adapted microorganisms to non-serpentine soils may constitute a biotic stressor that can induce changes in plant phenology.

Thirty-four genera across 5 phyla were shown to be differentially abundant between soil treatments by DESeq2 analysis. Of particular interest are *Solirubrobacter* which were only detected in nonserpentine soil treatments, *Lactobacillus* which characterized serpentine soil treatments, and *Methylobacterium* which were most abundant in NS+NSm, S+Sm, and NS+Sm treatments. *Solirubrobacter* are gram-positive, non-motile bacteria that has been identified and isolated from bulk soil, rhizosphere, and endosphere environments (Wei et al. 2014; Albuquerque and Da Costa 2014). In general, it has been shown to associate with high soil quality (Gravuer and Eskelinen 2017; Lopez et al. 2017; Sánchez-Marañón et al. 2017). Nonserpentine soils are generally more nutrient-rich than serpentine soils and this may influence the abundance of *Solirubrobacter* observed in the nonserpentine soil treatments. *Lactobacillus* are lactic acid bacteria (LAB) that are gram-positive and microaerophilic. Plasmids comprise up to 4.8% of LAB total gene content and are important for growth in the diverse, yet specific environments where these bacteria are found (Makarova et al. 2006). *Lactobacillus* have been shown to be metallotolerant and have the ability to bind heavy metals and protect against metal-induced oxidative stress (Li et al. 2017; Liu et al. 2019; Barman et al. 2020). Their abundance in serpentine soil treatments may reflect these phenotypic properties as serpentine soils have high concentrations of heavy metals. *Enterobacter*, which was most abundant in the NS+Sm+Drought treatment, has been shown to have plant-growth-promoting properties such as phosphorus solubility and ACC deaminase activity (Danish et al. 2020). Its application to *Sorghum bicolor* (L.) Moench increased the plants root architecture and ability to tolerate stress (Govindasamy et al. 2020). *Methylobacterium* (order *Rhizobiales*) belong to the same family as *Microvirga*, which has previously been shown to associate with legumes and non-legumes on serpentine soils (Igwe and Vannette 2019). In addition members of *Methylobacterium* can produce auxins and induce root nodulation (Kelly et al. 2014) and can promote plant growth through the production of ACC deaminase (Belimov et al. 2019; Sharma et al. 2021).

### Serpentine soils increase root diameter, but have no impact or decrease other plant growth metrics

*Plantago erecta* grown in serpentine soils were shorter, and generally smaller than those grown in nonserpentine soils as has been documented previously (O’Dell and Rajakaruna 2004; Kayama et al. 2005). Root length, surface area, and volume were smallest in serpentine soils while root diameter was the largest in this soil chemistry aligning with previous work that demonstrated that heavy-metal tolerant species of *Arabidopsis arenosa* and *Arabidopsis halleri* have thicker roots than the heavy-metal sensitive *Arabidopsis thaliana* (Staňová et al. 2012). Although all *P. erecta* growing in nonserpentine soils were larger than those in serpentine soils, only the plants in the NS+Sm soil treatment flowered sooner relative to the live serpentine (S+Sm) treatment. Collectively, differences in plant size and vegetative and flowering phenology between *P. erecta* on serpentine or nonserpentine soils are important for a plant’s life history (Metcalf et al. 2019). Plants that flower, set seed, and then die generally flower at the size that will ensure the best reproductive success (Metcalf et al. 2003). Continued research could determine how local adaptation of microbial communities influence reproductive success in plants growing in serpentine and nonserpentine soils.

Ecologically, competition for resources impact plant size. Soil nutrients, in particular, impact root architecture. Additionally, smaller plants are less vulnerable to drought (Olson et al. 2018). Phosphate deficiency, for example, produces plants that increase lateral root production over primary root production (López-Bucio et al. 2003). Plant-growth-promoting bacteria (PFPB) is one mechanism by which plants can access nutrients and defend against pathogenic bacteria (Glick 2012). A few direct plant-growth promoting methods that would be important in serpentine soils include phosphorus solubilization, metal chelation, and the production of extra-polymeric substances (EPS). Together, these traits would increase nutrient availability, decrease metal availability, and increase the water-holding capacity of the soil for the plant. Still, the ability of the microbes to confer benefits to plants growing on serpentine soils is dependent on local adaption of the microbes to serpentine (Rúa et al. 2016; Porter et al. 2016, 2019). Root diameter was the only plant trait that was larger is serpentine soils relative to nonserpentine soils and removing microbes that were locally adapted to serpentine soils removed this advantage. Conversely, replacing microbes that were locally adapted to nonserpentine with microbes that were locally adapted to serpentine contributed to plants that flowered sooner than other treatments.

## CONCLUSIONS

The root-associated microbial communities of *Plantago erecta* grown in serpentine and nonserpentine soils with adapted or non-adapted microorganisms have differing alpha and beta diversities. Notably, *P. erecta* grown in nonserpentine soils with serpentine microorganisms experienced accelerated vegetative and flowering phenology as they entered the vegetative, early flowering and late flowering phase before any plants that were grown in live serpentine soils (S+Sm) or live nonserpentine soil (NS+NSm). Above- and below-ground development on *P. erecta* on serpentine soil treatments were less than those grown on nonserpentine soil treatments. Overall, our results support a role of locally adapted microorganisms in impacting plant phenology despite minimal effects on other measurable aspects of plant phenotype.

## ACKNOWLEDGEMENTS

This work was made possible by the University of California Natural Reserve System (McLaughlin Natural Reserve) Reserve DOI: (https://doi.org/10.21973/N3W08D). We thank Cathy Koehler for assistance at McLaughlin Natural Reserve. We would like to thank Imade Ojo and Shenwen Gu who helped set up the experiment. Thanks to the California Native Plant Society, Davis Botanical Society Student Research Grant, Jastro Research Scholarship Award, and Natural Reserve System Graduate Student Grant Program for providing funds for the research. We are thankful to members of the Vannette Lab who provided feedback on manuscript drafts.

## REFERENCES

Akhtar MS, Siddiqui ZA, Wiemken A (2011) Arbuscular Mycorrhizal Fungi and Rhizobium to Control Plant Fungal Diseases. In: Lichtfouse E (ed) Alternative Farming Systems, Biotechnology, Drought Stress and Ecological Fertilisation. Springer Netherlands, Dordrecht, pp 263–292

Albuquerque L, Da Costa MS (2014) The families Conexibacteraceae, Patulibacteraceae and Solirubrobacteraceae. In: The Prokaryotes: Actinobacteria. Springer-Verlag Berlin Heidelberg, pp 185–200

Anacker BL (2014) The nature of serpentine endemism. Am J Bot 101:219–224. doi: 10.3732/ajb.1300349

Barillot CDC, Sarde C-O, Bert V, et al (2013) A standardized method for the sampling of rhizosphere and rhizoplan soil bacteria associated to a herbaceous root system. Ann Microbiol 63:471–476. doi: 10.1007/s13213-012-0491-y

Barman D, Jha DK, Bhattacharjee K (2020) Metallotolerant bacteria: Insights into bacteria thriving in metal-contaminated areas. In: Microbial Versatility in Varied Environments: Microbes in Sensitive Environments. plSpringer Singapore, pp 135–164

Belimov AA, Zinovkina NY, Safronova VI, et al (2019) Rhizobial ACC deaminase contributes to efficient symbiosis with pea (Pisum sativum L.) under single and combined cadmium and water deficit stress. Environ Exp Bot 167:103859. doi: 10.1016/j.envexpbot.2019.103859

Berg G, Smalla K (2009) Plant species and soil type cooperatively shape the structure and function of microbial communities in the rhizosphere. FEMS Microbiol Ecol 68:1–13. doi: 10.1111/j.1574-6941.2009.00654.x

Brady KU, Kruckeberg AR, Bradshaw Jr. HD (2005) Evolutionary ecology of plant adaptation to serpentine soils. Annu Rev Ecol Evol Syst 36:243–266. doi: 10.1146/annurev.ecolsys.35.021103.105730

Calderón K, Spor A, Breuil MC, et al (2017) Effectiveness of ecological rescue for altered soil microbial communities and functions. ISME J 11:272–283. doi: 10.1038/ismej.2016.86

Callahan BJ, McMurdie PJ, Rosen MJ, et al (2016a) DADA2: High-resolution sample inference from Illumina amplicon data. Nat Methods 13:581–583. doi: 10.1038/nmeth.3869

Callahan BJ, Sankaran K, Fukuyama JA, et al (2016b) Bioconductor workflow for microbiome data analysis: from raw reads to community analyses. F1000Research 5:1492. doi: 10.12688/f1000research.8986.2

Chaney L, Baucom RS (2020) The soil microbial community alters patterns of selection on flowering time and fitness-related traits in Ipomoea purpurea. Am J Bot 107:186–194. doi: 10.1002/ajb2.1426

Chaparro JM, Badri D V, Bakker MG, et al (2013) Root exudation of phytochemicals in Arabidopsis follows specific patterns that are developmentally programmed and correlate with soil microbial functions. PLoS One 8:e55731. doi: 10.1371/journal.pone.0055731

Chaparro JM, Badri D V, Vivanco JM (2014) Rhizosphere microbiome assemblage is affected by plant development. ISME J 8:790–803. doi: 10.1038/ismej.2013.196

Comeau AM, Li WKW, Tremblay JÉ, et al (2011) Arctic ocean microbial community structure before and after the 2007 record sea ice minimum. PLoS One 6:. doi: 10.1371/journal.pone.0027492

Cui QG, He WM (2009) Soil biota, but not soil nutrients, facilitate the invasion of Bidens pilosa relative to a native species Saussurea deltoidea. Weed Res 49:201–206. doi: 10.1111/j.1365-3180.2008.00679.x

Danish S, Zafar-Ul-Hye M, Hussain S, et al (2020) Mitigation of drought stress in maize through inoculation with drought tolerant ACC deaminase containing PGPR under axenic conditions. Pakistan J Bot 52:49–60. doi: 10.30848/PJB2020-1(7)

De Curtis F, Lima G, Vitullo D, De Cicco V (2010) Biocontrol of Rhizoctonia solani and Sclerotium rolfsii on tomato by delivering antagonistic bacteria through a drip irrigation system. Crop Prot 29:663–670. doi: 10.1016/j.cropro.2010.01.012

Dombrowski N, Schlaeppi K, Agler MT, et al (2017) Root microbiota dynamics of perennial Arabis alpina are dependent on soil residence time but independent of flowering time. ISME J 11:43–55. doi: 10.1038/ismej.2016.109

Dorji T, Totland Ø, Moe SR, et al (2013) Plant functional traits mediate reproductive phenology and success in response to experimental warming and snow addition in Tibet. Glob Chang Biol 19:459–472. doi: 10.1111/gcb.12059

Edwards JA, Santos-Medellín CM, Liechty ZS, et al (2018) Compositional shifts in root-associated bacterial and archaeal microbiota track the plant life cycle in field-grown rice. PLoS Biol 16:. doi: 10.1371/journal.pbio.2003862

Emami S, Alikhani HA, Pourbabaei AA, et al (2018) Improved growth and nutrient acquisition of wheat genotypes in phosphorus deficient soils by plant growth-promoting rhizospheric and endophytic bacteria. Soil Sci Plant Nutr 64:719–727. doi: 10.1080/00380768.2018.1510284

Farrer EC, Suding KN (2016) Teasing apart plant community responses to N enrichment: the roles of resource limitation, competition and soil microbes. Ecol Lett 19:1287–1296. doi: 10.1111/ele.12665

Fei H, Crouse M, Papadopoulos YA, Vessey JK (2020) Improving biomass yield of giant Miscanthus by application of beneficial soil microbes and a plant biostimulant. Can J Plant Sci 100:29–39. doi: 10.1139/cjps-2019-0012

Fenner M (1998) The phenology of growth and reproduction in plants. Perspect Plant Ecol Evol Syst 1:78–91. doi: https://doi.org/10.1078/1433-8319-00053

Glick BR (2012) Plant Growth-Promoting Bacteria: Mechanisms and Applications. 963401:. doi: 10.6064/2012/963401

Glöckner FO, Yilmaz P, Quast C, et al (2017) 25 years of serving the community with ribosomal RNA gene reference databases and tools. J Biotechnol 261:169–176. doi: 10.1016/j.jbiotec.2017.06.1198

Gouda S, Kerry RG, Das G, et al (2018) Revitalization of plant growth promoting rhizobacteria for sustainable development in agriculture. Microbiol Res 206:131–140. doi: 10.1016/j.micres.2017.08.016

Govindasamy V, George P, Kumar M, et al (2020) Multi-trait PGP rhizobacterial endophytes alleviate drought stress in a senescent genotype of sorghum [Sorghum bicolor (L.) Moench]. 3 Biotech 10:13. doi: 10.1007/s13205-019-2001-4

Gravuer K, Eskelinen A (2017) Nutrient and rainfall additions shift phylogenetically estimated traits of soil microbial communities. Front Microbiol 8:1–16. doi: 10.3389/fmicb.2017.01271

Haichar FEZ, Marol C, Berge O, et al (2008) Plant host habitat and root exudates shape soil bacterial community structure. ISME J 2:1221–1230. doi: 10.1038/ismej.2008.80

Hidalgo-Triana N, Pérez-Latorre A V. (2018) Phenological patterns in Mediterranean south Iberian serpentine flora. Nord J Bot 36:1–11. doi: 10.1111/njb.02028

Igwe AN, Vannette RL (2019) Bacterial communities differ between plant species and soil type, and differentially influence seedling establishment on serpentine soils. Plant Soil 441:423– 437. doi: 10.1007/s11104-019-04135-5

Kaisermann A, de Vries FT, Griffiths RI, Bardgett RD (2017) Legacy effects of drought on plant–soil feedbacks and plant–plant interactions. New Phytol 215:1413–1424. doi: 10.1111/nph.14661

Kandlikar GS, Johnson CA, Yan X, et al (2019) Winning and losing with microbes: how microbially mediated fitness differences influence plant diversity. Ecol Lett 22:1178–1191. doi: 10.1111/ele.13280

Kassambara A, Kosinski M (2018) survminer: Drawing Survival Curves using “ggplot2”. R package version 0.4.3

Kayama M, Quoreshi AM, Uemura S, Koike T (2005) Differences in growth characteristics and dynamics of elements absorbed in seedlings of three spruce species raised on serpentine soil in northern Japan. Ann Bot 95:661–672. doi: 10.1093/aob/mci063

Kazan K, Lyons R (2016) The link between flowering time and stress tolerance. J Exp Bot 67:47–60. doi: 10.1093/jxb/erv441

Kelly DP, McDonald IR, Wood AP (2014) The Family Methylobacteriaceae. In: Rosenberg E, DeLong EF, Lory S, et al. (eds) The Prokaryotes. Spring, Berlin, Heidelberg, pp 313–340

Kwak MJ, Kong HG, Choi K, et al (2018) Rhizosphere microbiome structure alters to enable wilt resistance in tomato. Nat Biotechnol 36:1100–1116. doi: 10.1038/nbt.4232

Lau JA, Lennon JT (2012) Rapid responses of soil microorganisms improve plant fitness in novel environments. Proc Natl Acad Sci 109:14058–14062. doi: 10.1073/pnas.1202319109

Li B, Jin D, Yu S, et al (2017) In Vitro and in Vivo evaluation of Lactobacillus delbrueckii subsp. Bulgaricus KLDS1.0207 for the alleviative effect on lead toxicity. Nutrients 9:p. doi: 10.3390/nu9080845

Liu, Zheng, Ma, et al (2019) Evaluation and Proteomic Analysis of Lead Adsorption by Lactic Acid Bacteria. Int J Mol Sci 20:5540. doi: 10.3390/ijms20225540

López-Bucio J, Cruz-Ramírez A, Herrera-Estrella L (2003) The role of nutrient availability in regulating root architecture. Curr Opin Plant Biol 6:280–287. doi: 10.1016/S1369-5266(03)00035-9

Lopez S, Piutti S, Vallance J, et al (2017) Nickel drives bacterial community diversity in the rhizosphere of the hyperaccumulator Alyssum murale. Soil Biol Biochem 114:121–130. doi: 10.1016/j.soilbio.2017.07.010

Lu T, Ke M, Lavoie M, et al (2018) Rhizosphere microorganisms can influence the timing of plant flowering. Microbiome 6:. doi: 10.1186/s40168-018-0615-0

Makarova K, Slesarev A, Wolf Y, et al (2006) Comparative genomics of the lactic acid bacteria. Proc Natl Acad Sci U S A 103:15611–15616. doi: 10.1073/pnas.0607117103

McMurdie PJ, Holmes S (2013) Phyloseq: An R Package for Reproducible Interactive Analysis and Graphics of Microbiome Census Data. PLoS One 8:p. doi: 10.1371/journal.pone.0061217

Metcalf CJE, Henry LP, Rebolleda-Gómez M, Koskella B (2019) Why evolve reliance on the microbiome for timing of ontogeny? MBio 10:1–10. doi: 10.1128/mBio.01496-19

Metcalf JC, Rose KE, Rees M (2003) Evolutionary demography of monocarpic perennials. Trends Ecol Evol 18:471–480. doi: 10.1016/S0169-5347(03)00162-9

Murgese P, Santamaria P, Leoni B, Crecchio C (2020) Ameliorative Effects of PGPB on Yield, Physiological Parameters, and Nutrient Transporter Genes Expression in Barattiere (Cucumis melo L.). J Soil Sci Plant Nutr 20:784–793. doi: 10.1007/s42729-019-00165-1

O’Dell RE, James JJ, Richards JH (2006) Congeneric serpentine and nonserpentine shrubs differ more in leaf Ca:Mg than in tolerance of low N, low P, or heavy metals. Plant Soil 280:49– 64. doi: 10.1007/s11104-005-3502-y

O’Dell RE, Rajakaruna N (2004) Intraspecific Variation, Adaptation, and Evolution. In: ed Harrison S, Rajakaruna N (eds) Serpentine: The Evolution and Ecology of a Model System. pp 97– 138

Oksanen J, Blanchet FG, Friendly M, et al (2019) Vegan: Community Ecology Package (version 2.5-6)

Oline DK (2006) Phylogenetic comparisons of bacterial communities from serpentine and nonserpentine soils. Appl Environ Microbiol 72:6965–6971. doi: 10.1128/AEM.00690-06

Olson ME, Soriano D, Rosell JA, et al (2018) Plant height and hydraulic vulnerability to drought and cold. Proc Natl Acad Sci U S A 115:7551–7556. doi: 10.1073/pnas.1721728115

Orlandini V, Emiliani G, Fondi M, et al (2014) Network Analysis of Plasmidomes: The Azospirillum brasilense Sp245 Case . Int J Evol Biol 2014:1–14. doi: 10.1155/2014/951035

Paterson E, Gebbing T, Abel C, et al (2007) Rhizodeposition shapes rhizosphere microbial community structure in organic soil. New Phytol 173:600–610. doi: 10.1111/j.1469-8137.2006.01931.x

Porter SS, Bantay R, Friel CA, et al (2019) Beneficial microbes ameliorate abiotic and biotic sources of stress on plants. doi: 10.1111/1365-2435.13499

Porter SS, Chang PL, Conow CA, et al (2017) Association mapping reveals novel serpentine adaptation gene clusters in a population of symbiotic Mesorhizobium. ISME J 11:248–262. doi: 10.1038/ismej.2016.88

Porter SS, Chang PL, Conow CA, et al (2016) Association mapping reveals novel serpentine adaptation gene clusters in a population of symbiotic Mesorhizobium. Isme J 11:248

Quast C, Pruesse E, Yilmaz P, et al (2012) The SILVA ribosomal RNA gene database project: improved data processing and web-based tools. Nucleic Acids Res 41:D590–D596. doi: 10.1093/nar/gks1219

Ravichandran KR, Thangavelu M (2017) Role and influence of soil microbial communities on plant invasion. Ecol Quest 27:9–23. doi: 10.12775/EQ.2017.024

Rodríguez-Pérez J, Traveset A (2016) Effects of flowering phenology and synchrony on the reproductive success of a long-flowering shrub. AoB Plants 8:p. doi: 10.1093/aobpla/plw007

Rossington N, Yost J, Ritter M (2018) Water Availability Influences Species Distributions on Serpentine Soils. Madroño 65:68–79. doi: 10.3120/0024-9637-65.2.68

Rúa MA, Antoninka A, Antunes PM, et al (2016) Home-field advantage? evidence of local adaptation among plants, soil, and arbuscular mycorrhizal fungi through meta-analysis. BMC Evol Biol 16:1–15. doi: 10.1186/s12862-016-0698-9

Safford AHD, Viers JH, Harrison SP (2005) Serpentine endemism in the California flora: a database of serpentine affinity. Madrono 52:222–257

Sakaguchi S, Horie K, Ishikawa N, et al (2019) Maintenance of soil ecotypes of Solidago virgaurea in close parapatry via divergent flowering time and selection against immigrants. J Ecol 107:418–435. doi: 10.1111/1365-2745.13034

Sánchez-Marañón M, Miralles I, Aguirre-Garrido JF, et al (2017) Changes in the soil bacterial community along a pedogenic gradient. Sci Rep 7:1–11. doi: 10.1038/s41598-017-15133-x

Schliep KP (2011) phangorn: Phylogenetic analysis in R. Bioinformatics 27:592–593. doi: 10.1093/bioinformatics/btq706

Schneider A (2017) Flowering time evolution is independent of serpentine tolerance in the California flora. Ecosphere 8:. doi: 10.1002/ecs2.1767

Sharma S, Chandra D, Sharma AK (2021) Rhizosphere Plant–Microbe Interactions Under Abiotic Stress. Springer, Singapore, pp 195–216

Sherrard ME, Maherali H (2006) the Adaptive Significance of Drought Escape in Avena Barbata, an Annual Grass. Evolution (N Y) 60:2478. doi: 10.1554/06-150.1

Staňová A, Ďurišová E, Banásová V, et al (2012) Root system morphology and primary root anatomy in natural non-metallicolous and metallicolous populations of three Arabidopsis species differing in heavy metal tolerance. Biologia (Bratisl) 67:505–516. doi: 10.2478/s11756-012-0040-y

Therneau TM (2015) A Package for Survival Analysis in S. version 2.38

Van Der Heijden MGA, Bakker R, Verwaal J, et al (2006) Symbiotic bacteria as a determinant of plant community structure and plant productivity in dune grassland. FEMS Microbiol Ecol 56:178–187. doi: 10.1111/j.1574-6941.2006.00086.x

Wagner MR, Lundberg DS, Coleman-Derr D, et al (2014) Natural soil microbes alter flowering phenology and the intensity of selection on flowering time in a wild Arabidopsis relative. Ecol Lett 17:717–726. doi: 10.1111/ele.12276

Wei L, Ouyang S, Wang Y, et al (2014) Solirubrobacter phytolaccae sp. nov., an endophytic bacterium isolated from roots of Phytolacca acinosa Roxb. Int J Syst Evol Microbiol 64:858–862. doi: 10.1099/ijs.0.057554-0

Wright JW, Stanton ML, Scherson R (2006) Local adaptation to serpentine and non-serpentine soils in Collinsia sparsiflora. Evol Ecol Res 8:1–21

Yang X, Guo R, Knops JMH, et al (2020) Shifts in plant phenology induced by environmental changes are small relative to annual phenological variation. Agric For Meteorol 294:108144. doi: 10.1016/j.agrformet.2020.108144

Yilmaz P, Parfrey LW, Yarza P, et al (2014) The SILVA and “all-species Living Tree Project (LTP)” taxonomic frameworks. Nucleic Acids Res 42:643–648. doi: 10.1093/nar/gkt1209

Zhalnina K, Louie KB, Hao Z, et al (2018) Dynamic root exudate chemistry and microbial substrate preferences drive patterns in rhizosphere microbial community assembly. Nat Microbiol 3:470–480. doi: 10.1038/s41564-018-0129-3

