## Supplemental Figures for "Plant phenology influences rhizosphere microbial community and is accelerated by serpentine microorganisms in *Plantago erecta*"

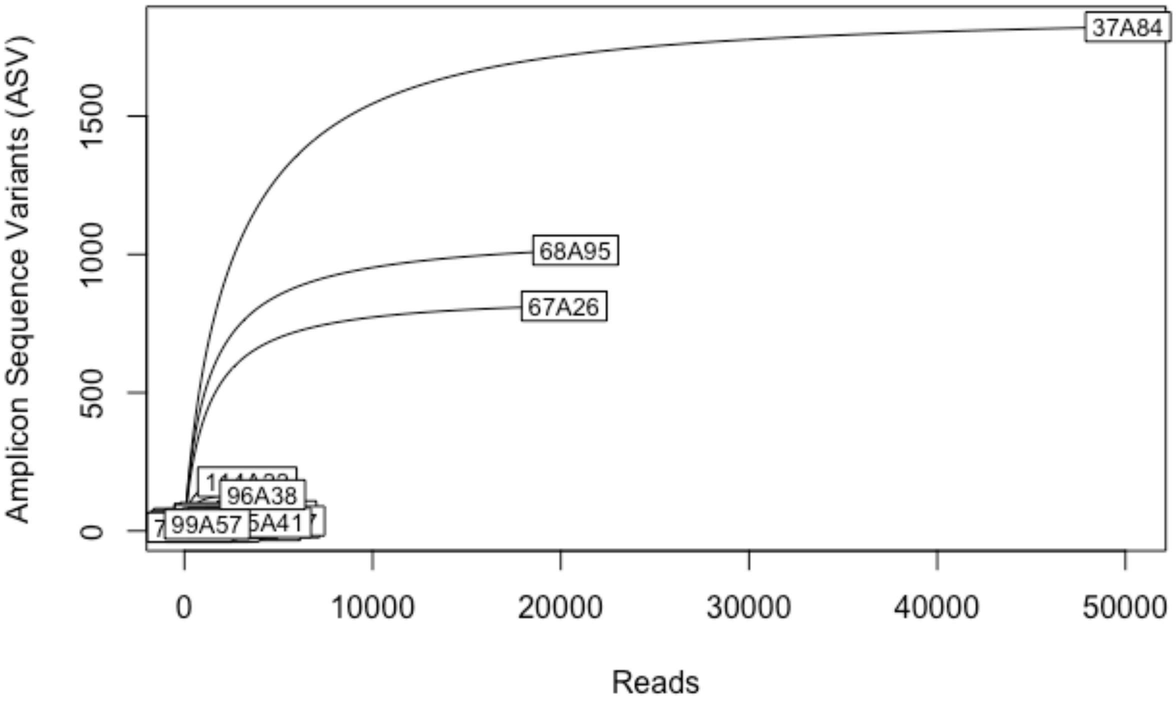


Supplemental Figure 1 - Sampling curves to 50,000 sequences for cumulative reads from the Serpentine soil and Serpentine microbes (S+Sm) treatment (A).

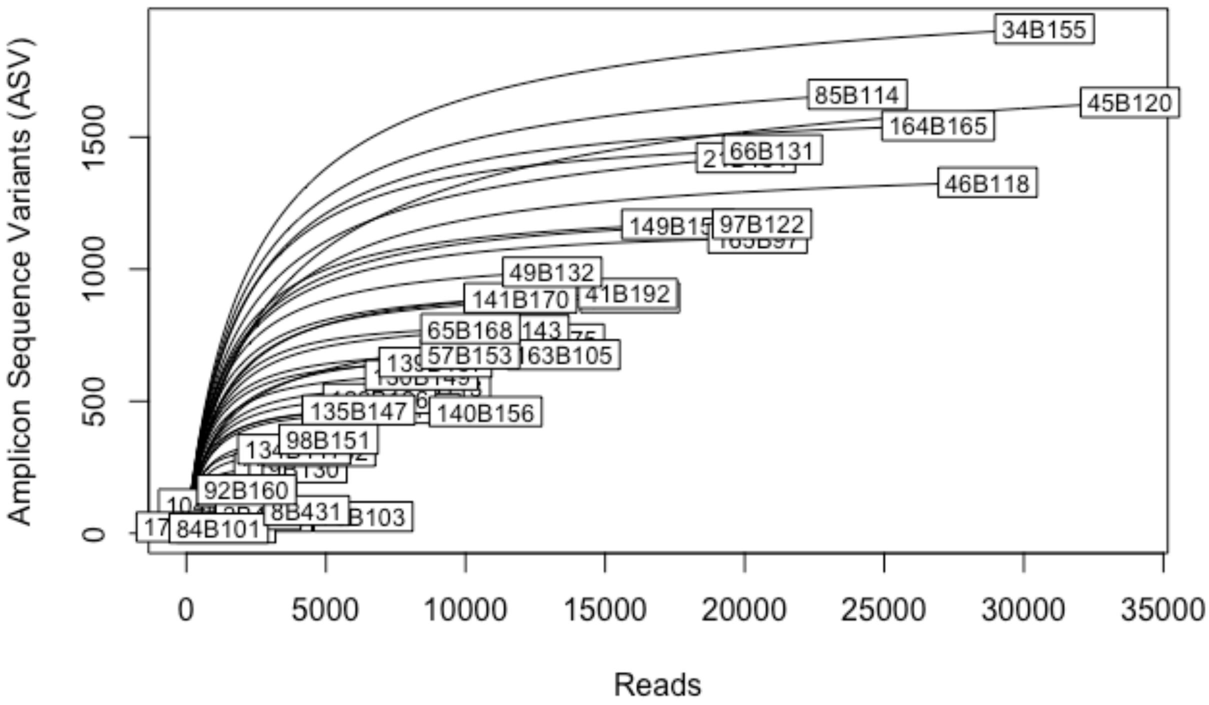


Supplemental Figure 2 - Sampling curves to 35,000 sequences for cumulative reads from the Nonserpentine soil and Nonserpentine microbes (NS and NSm) treatment (B).

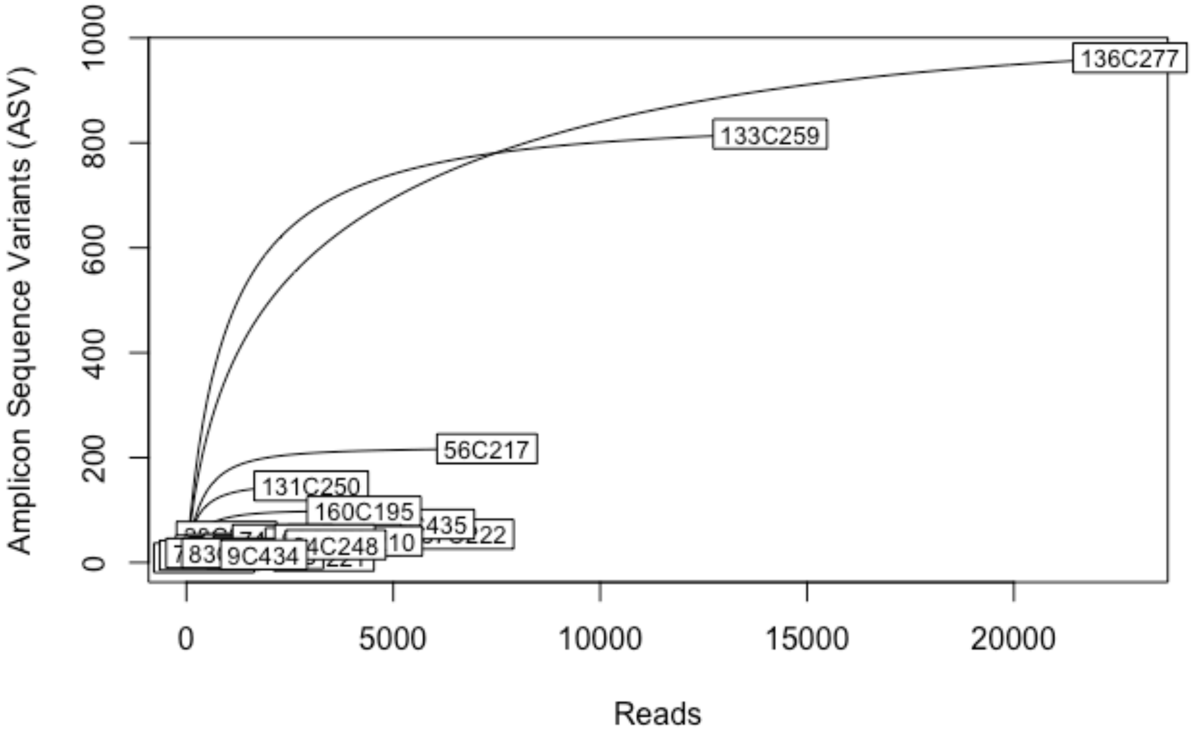


Supplemental Figure 3 - Sampling curves to 25,000 sequences for cumulative reads from the Nonserpentine soil and Serpentine microbes (NS and Sm) treatment (C).

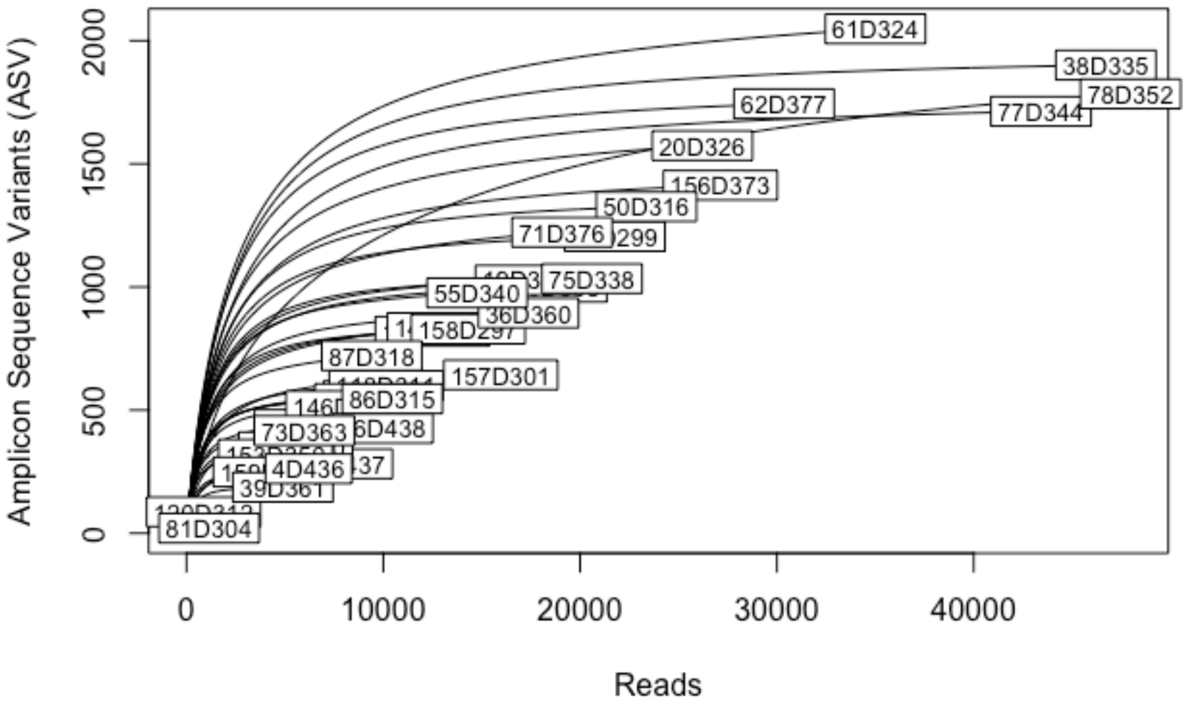


Supplemental Figure 4 - Sampling curves to 45,000 sequences for cumulative reads from the Serpentine soil and Nonserpentine microbe (S and NSm) treatment (D).

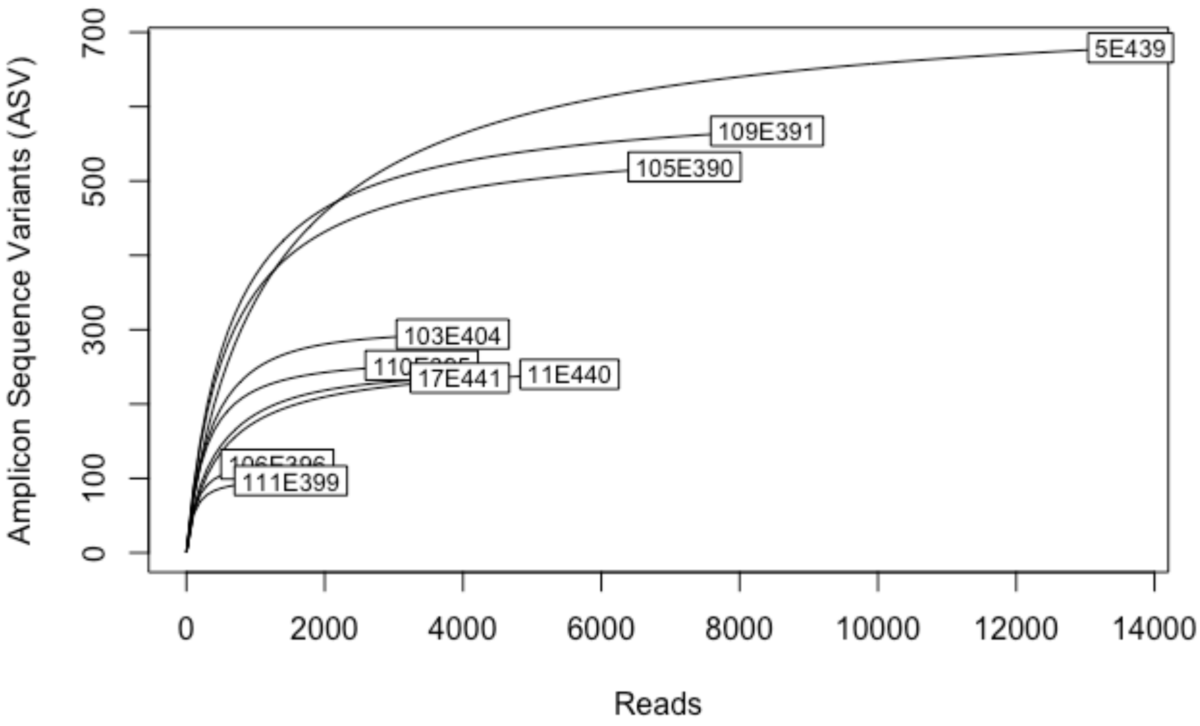


Supplemental Figure 5 - Sampling curves to 14,000 sequences for cumulative reads from the Nonserpentine soil and Serpentine microbe and Nickel stress (NS and Sm and Ni) treatment (E).

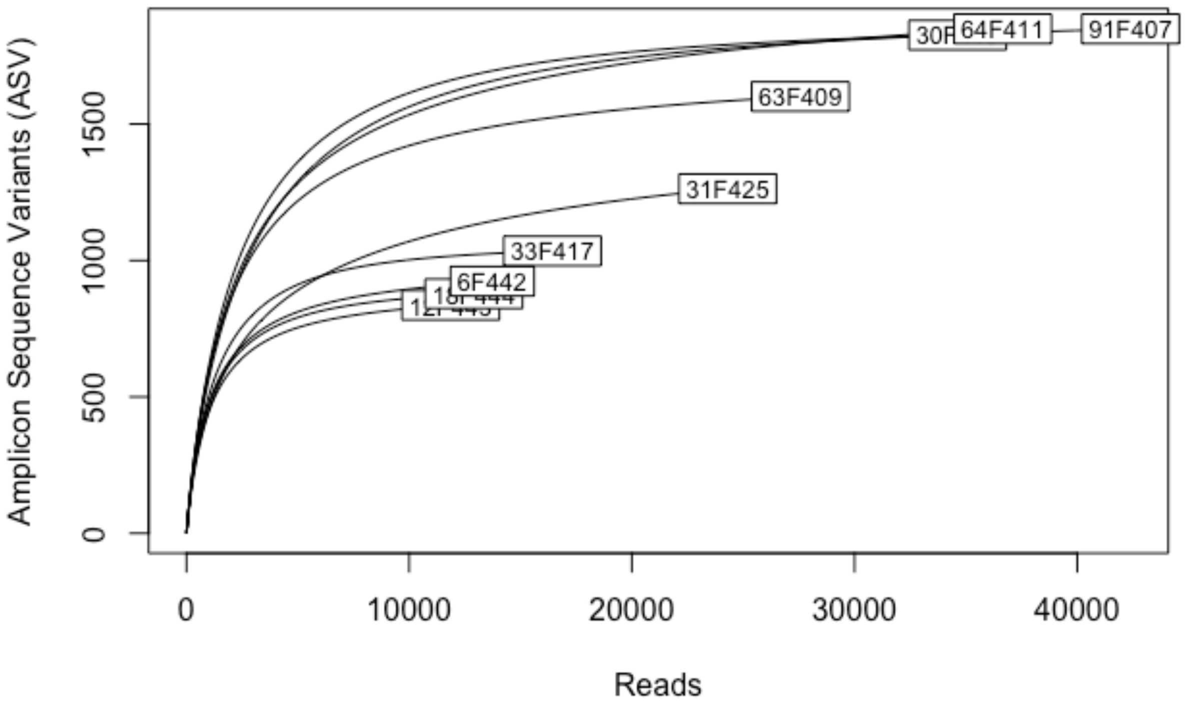


Supplemental Figure 6 - Sampling curves to 45,000 sequences for cumulative reads from the Nonserpentine and Serpentine microbe and Drought stress (NS and Sm and Drought) treatment (F).

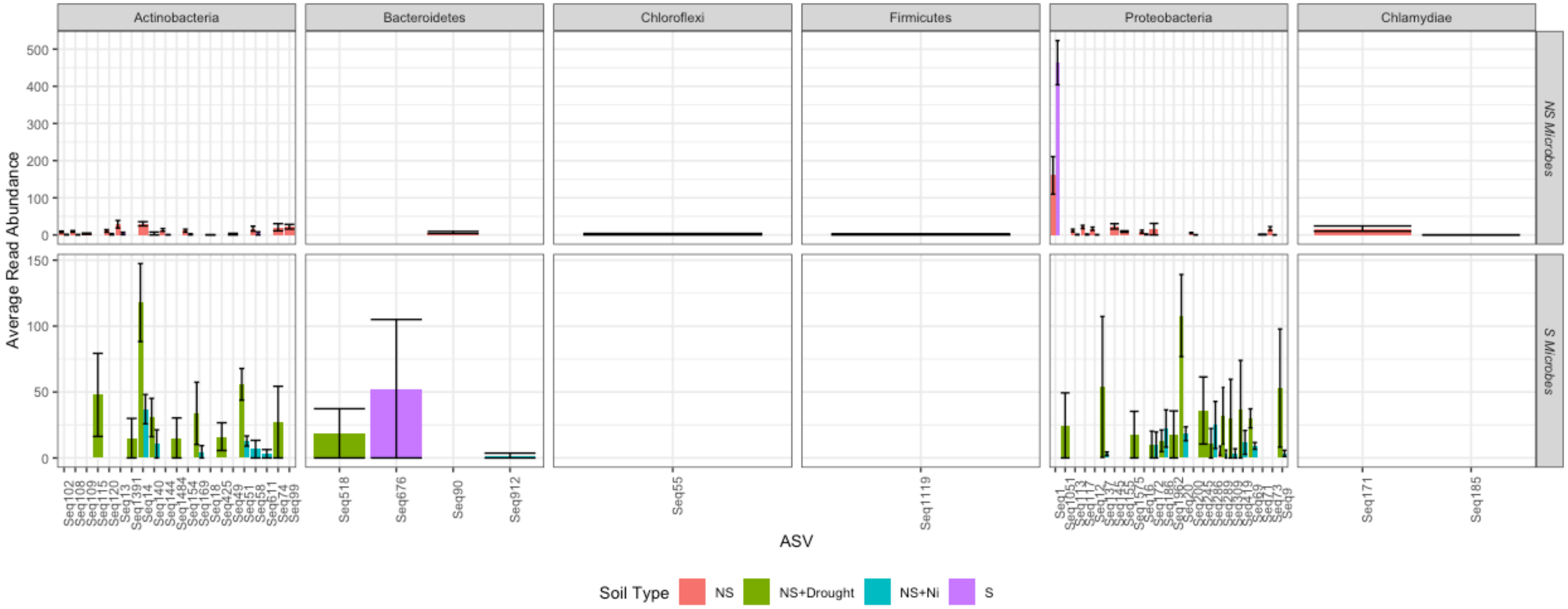


Supplemental Figure 7 - Differentially abundant ASV between soil types based on DESeq2 analysis (FDR <0.01) faceted by genus and microbe source with ASV on the x-axis. Each color represents a distinct soil treatment. Bars represent means +/- 1SE.

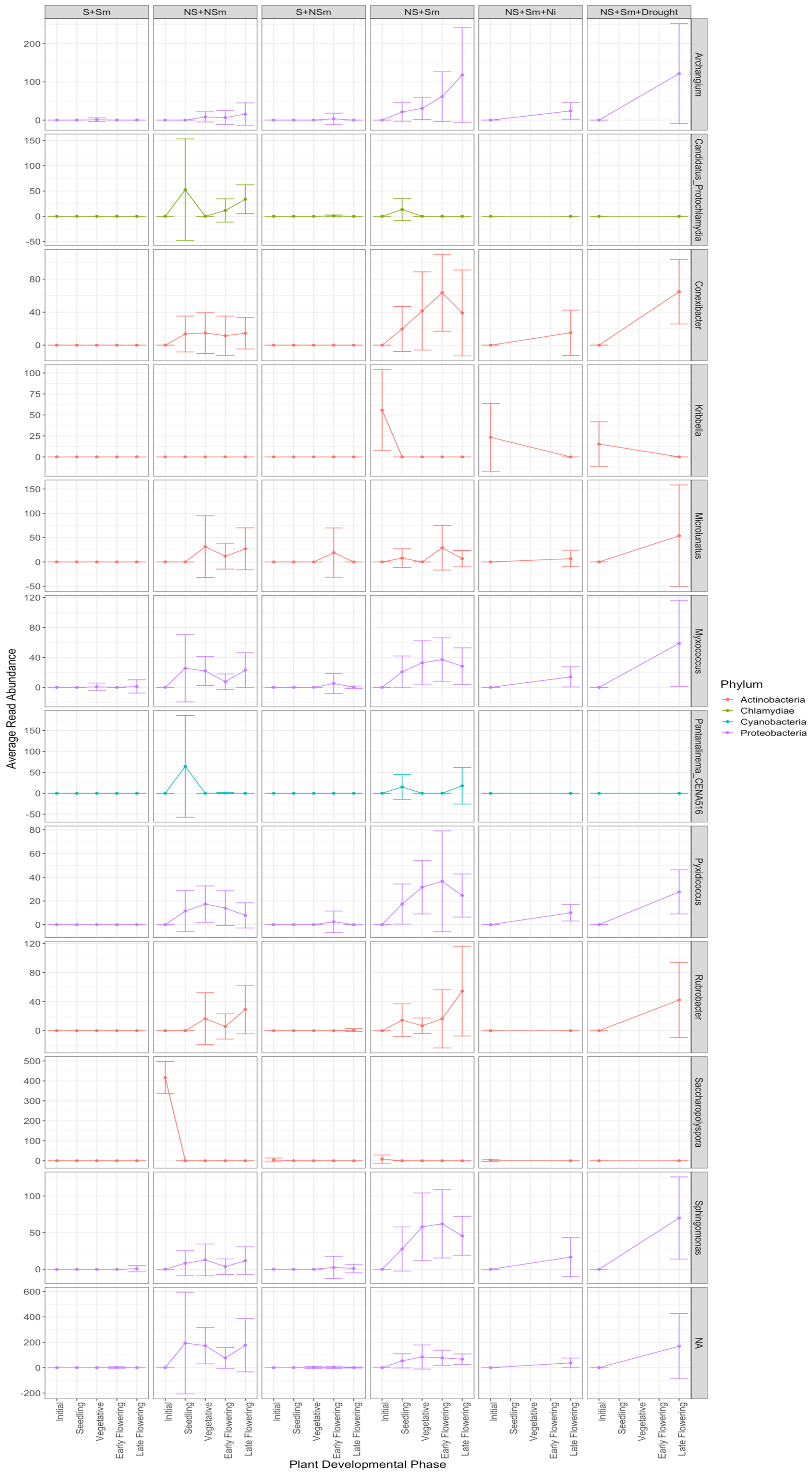


Supplemental Figure 8 - Differentially abundant ASV based on DESeq2 analysis (FDR <0.01) between plant developmental phases facetted by genus with each color representing a distinct phylum. Points represent means +/- 1SE.
